## Supplement for "Predicting the locations of cryptic pockets from single protein structures using the PocketMiner graph neural network"

### Supplementary information

#### Supplementary figures

Contents

1. Featurization of TEM-1 β-lactamase
2. Label consistency in independent simulations
3. RMSD distributions
4. Ligands of carbonic anhydrase
5. Curation of negative examples
6. Structural alignments of high sequence identity proteins
7. PocketMiner predictions on the test set
8. PocketMiner predictions for Wnt2 and simulations of Wnt2
9. 3D CNN grid featurization
10. 3D CNN training and validation PR-AUC

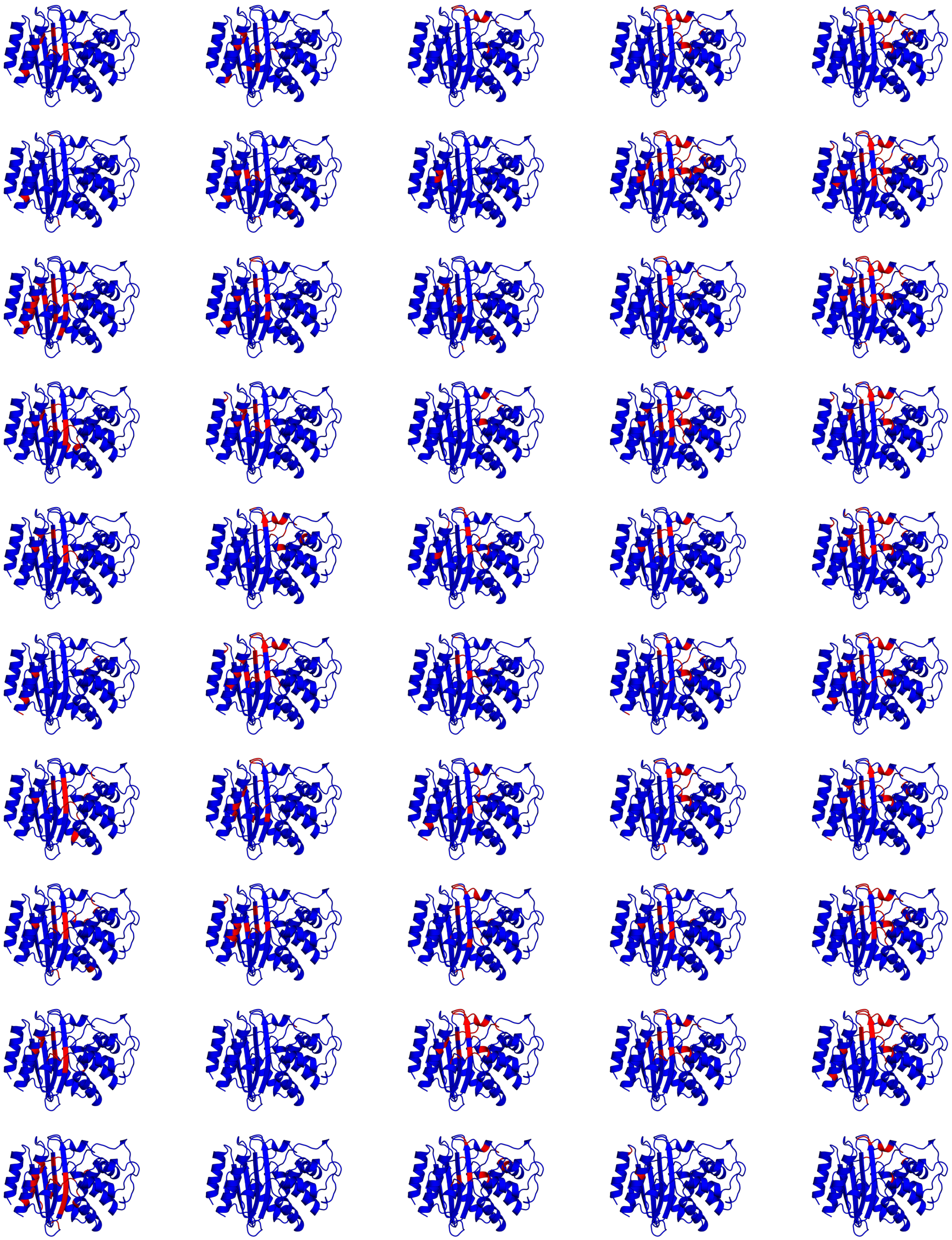

**Figure S1:** Featurization of TEM-1 β-lactamase. Featurization repeatedly identifies residues of both the horn site and omega loop pockets as cryptic sites without identifying residues in most other areas of the protein. Residues with negative featurization labels are colored blue while those with positive labels are colored red. Labels from 5 rounds (columns) of FAST with 10 parallel simulations each (rows) are shown projected onto the *apo* crystal structure used as a starting structure, with subsequent rounds running from left to right.

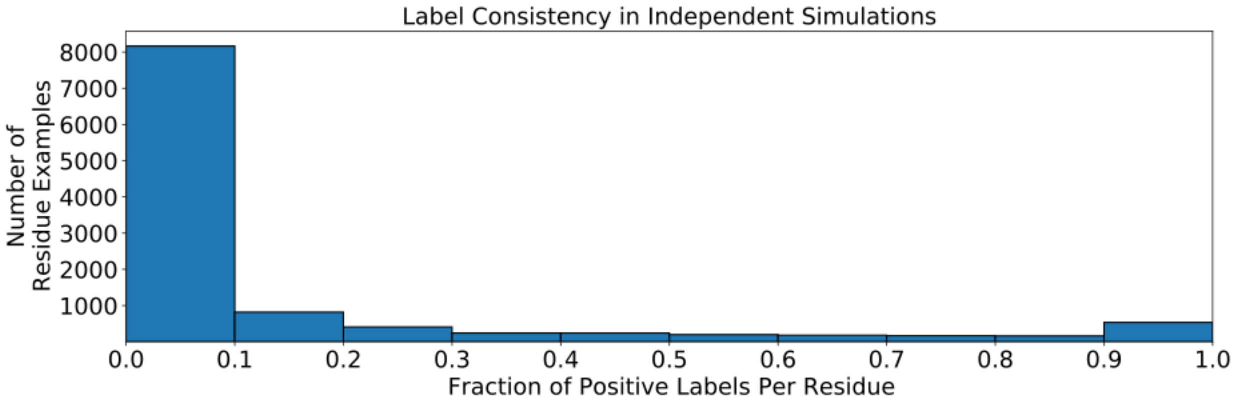


**Figure S2:** Label consistency in independent simulations. We determined the fraction of times that each residue participates in a cryptic pocket opening across independent 40 ns-long simulations launched from the same starting structure. Values of 0 and 1 indicate perfect consistency (either opening in no simulations or opening in all simulations). The two peaks at 0 and 1 indicate that our labeling scheme produces consistent labels across independent simulations launched from the same structure. However, among positive labels, a positive example was more likely to only open in a minority of independent simulations, indicating that many pocket opening events are rare on the timescales analyzed (40 ns).

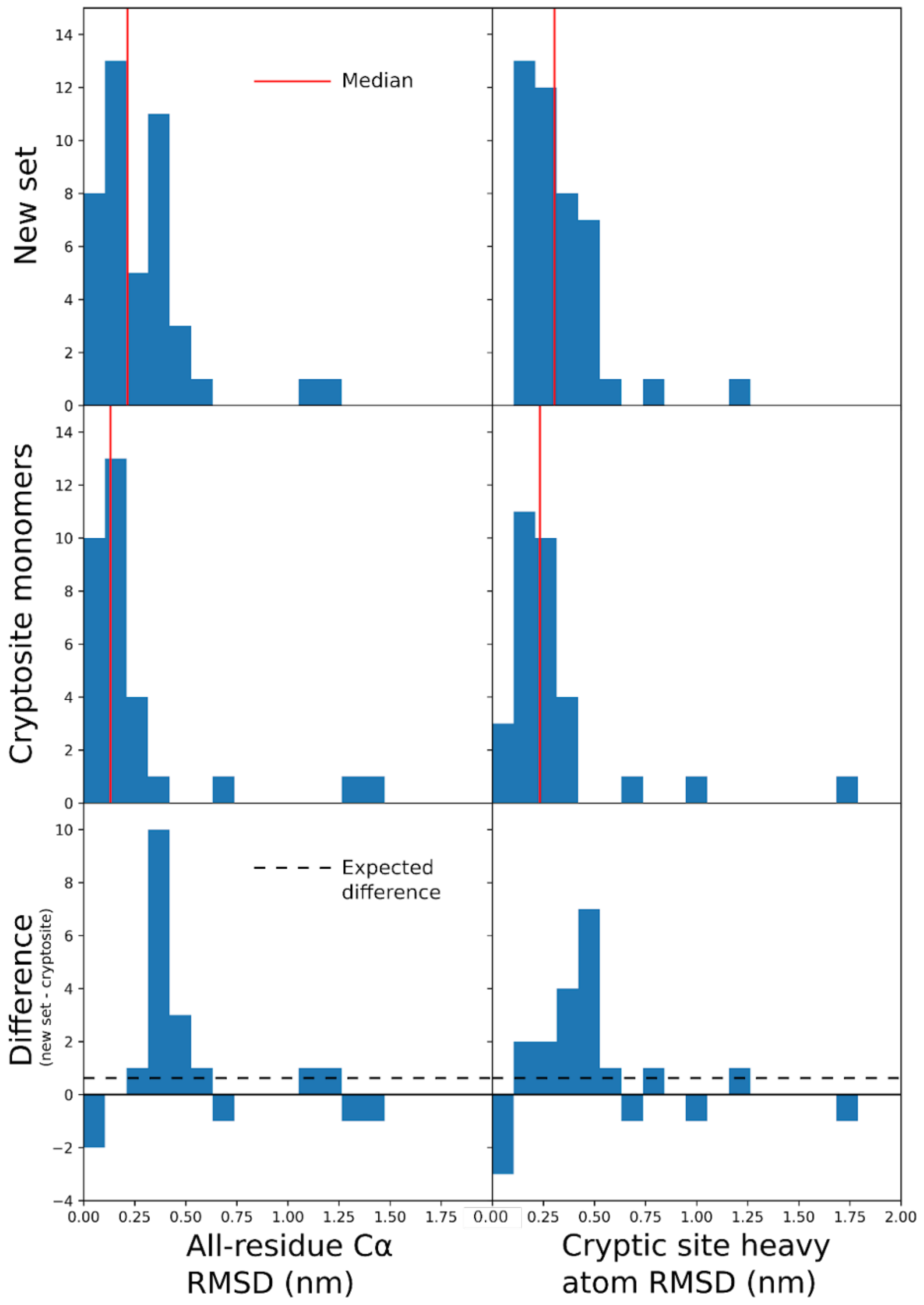


**Figure S3:** RMSD distributions for the new protein set compiled in this work and for physiological monomers in the CryptoSite set without large gaps. Medians are shown in red.

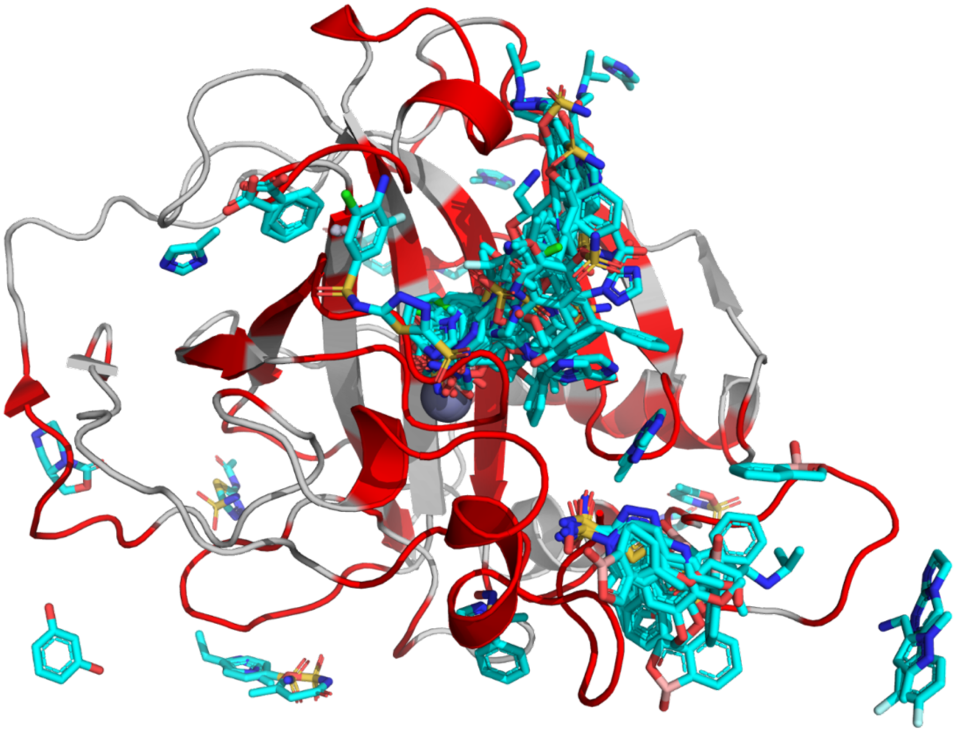

**Figure S4:** Ligands binding carbonic anhydrase. The carbonic anhydrase cluster centroid (PDB ID 1YDA) is shown as a ribbon. Residues which are within 5Å of a valid MOAD ligand in any centroid are shown in red, while the remainder, which were used as negative true labels for testing and validation, are shown in white. A subset of ligands which are collectively within 5Å of every nonnegative residue are shown as cyan sticks.

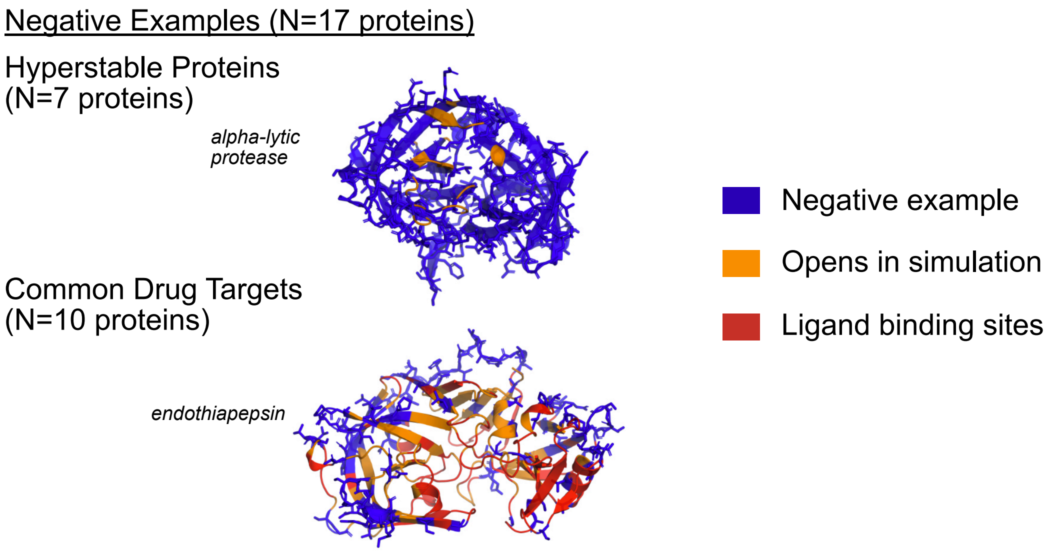


**Figure S5:** Negative examples were curated from hyper-rigid proteins and common drug targets. We ran simulations of both kinds of proteins to determine if any pockets formed during these simulations. Any residues that were adjacent to a pocket in simulation were excluded from the test set. For common drug targets, any residues that bind a ligand were also excluded from the test set.

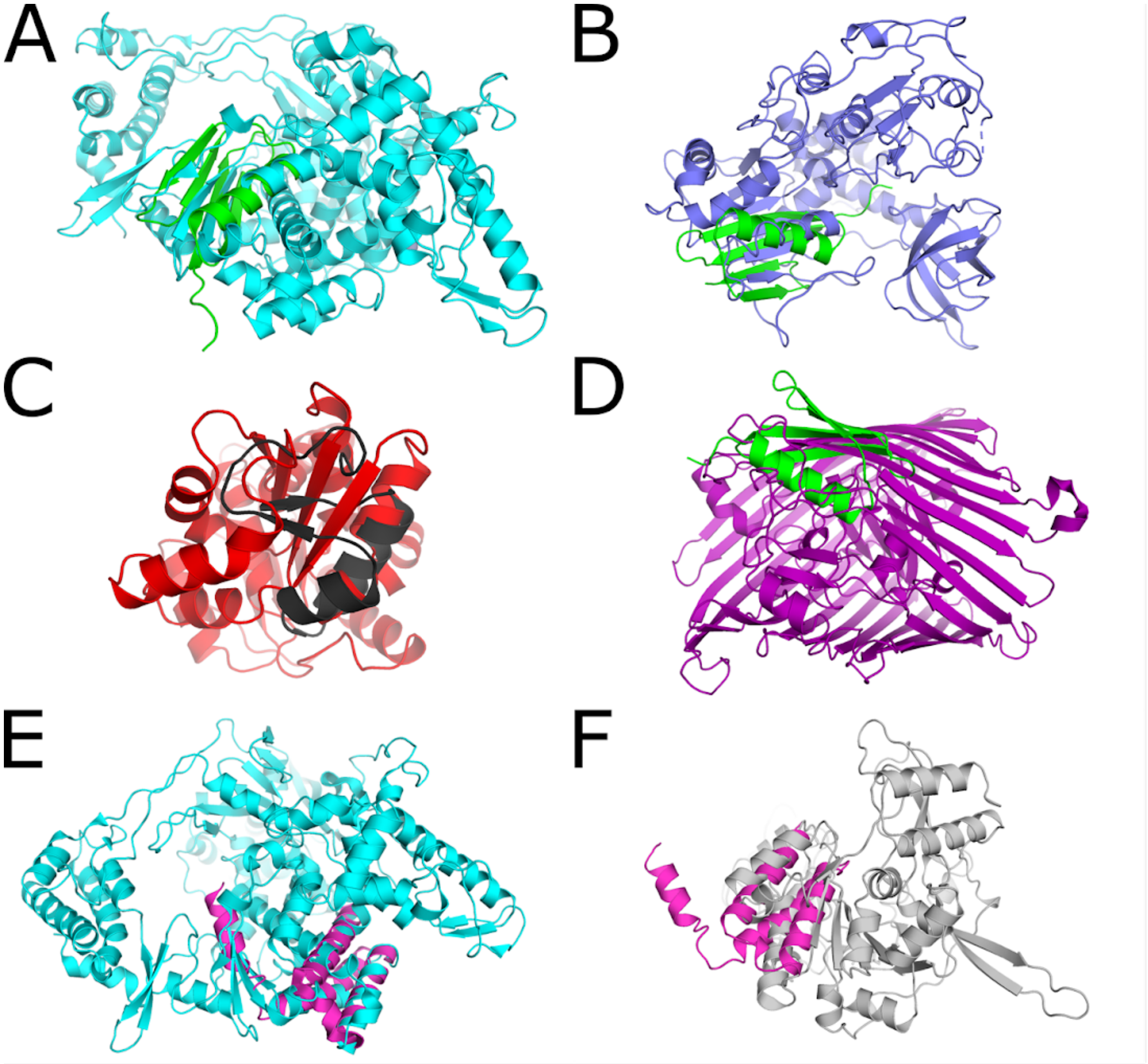


**Figure S6:** Structural alignments of high-sequence-identity pairs of proteins used in this paper, using PyMol's *cealign* function (see table S2).

A. SARS-CoV-2 nsp12 (cyan) and PDB ID 1IGD (green). 1IGD is ~1/10$^{th}$ the length of nsp12. It aligns reasonably well to one area of nsp12, but the beta strand polarity and topology of the loops connecting the beta strands does not match.

B. PDB ID 1IGD and SARS-CoV-2 nsp13 (blue). Once again, 1IGD aligns reasonably well to one area of the much larger nsp13, but the beta strand polarity and topology of the loops connecting the beta strands does not match.

C. PDB ID 2FD7 (black) and PDB ID 2OHG (red), showing no meaningful structural homology.

D. PDB ID 1IGD and PDB ID 1KMO (purple), showing no meaningful structural homology.

E. SARS-CoV-2 nsp12 and SARS-CoV-2 nsp7 (magenta), showing no meaningful structural homology.

F. PDB ID 5NZM (grey) and SARS-CoV-2 nsp7, showing no meaningful structural homology.

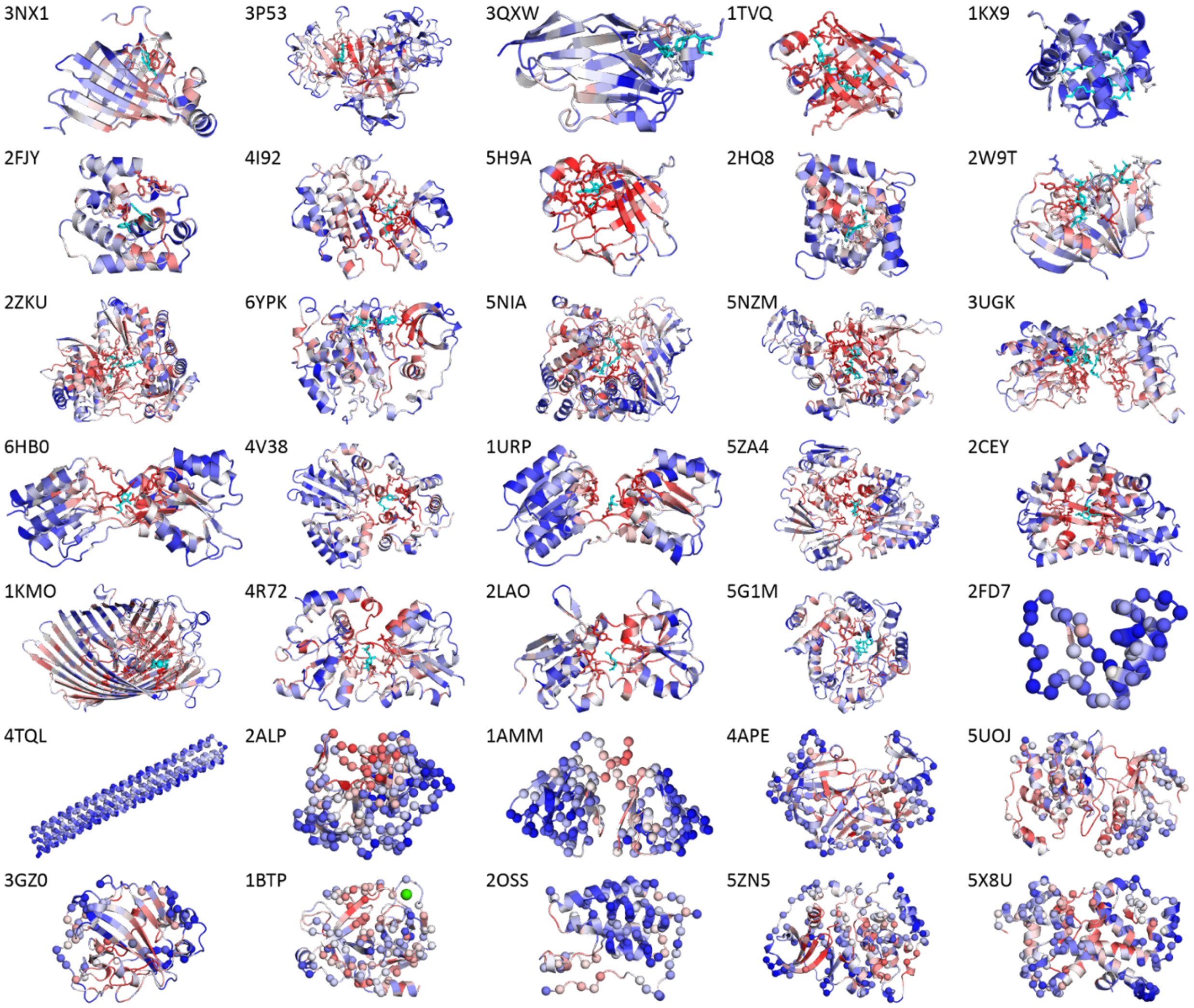


**Figure S7:** PocketMiner predictions on the test set. Proteins are shown as ribbons, colored from blue to red as predictions range from negative to positive. Ligand-lining residues (positive true labels) are shown as sticks. Residues in proteins believed to lack cryptic pockets which did not line pockets in simulation and residues in well-studied proteins which were neither adjacent to drug-like ligands nor lined pockets in simulations (negative true labels) are shown in spheres. Protein-bound ions are shown as spheres and colored according to their respective elements. Ligands binding in cryptic pockets are shown in cyan sticks (and cyan spheres for the two iron ions which comprise part of PDB ID 1KMO's cryptic ligand assembly). Proteins are ordered left to right and then top to bottom, with cryptic pocket examples first in order of decreasing difference between the mean distance between ligand-lining residues in *holo* and *apo* (see attached SI spreadsheet tab validation_and_test_sets, columns O-R). Forward pockets are listed first, followed by ones involving a mixture of forward and reverse motions, followed by reverse pockets. 2FJY is listed out of order and placed next to the functionally related protein 1KX9 because the large number of *apo* residues which become unresolved in *holo* render the assignment of pocket direction unreliable. After the cryptic pocket examples are three proteins believed to be highly rigid, followed by proteins with many *holo* crystal structures.

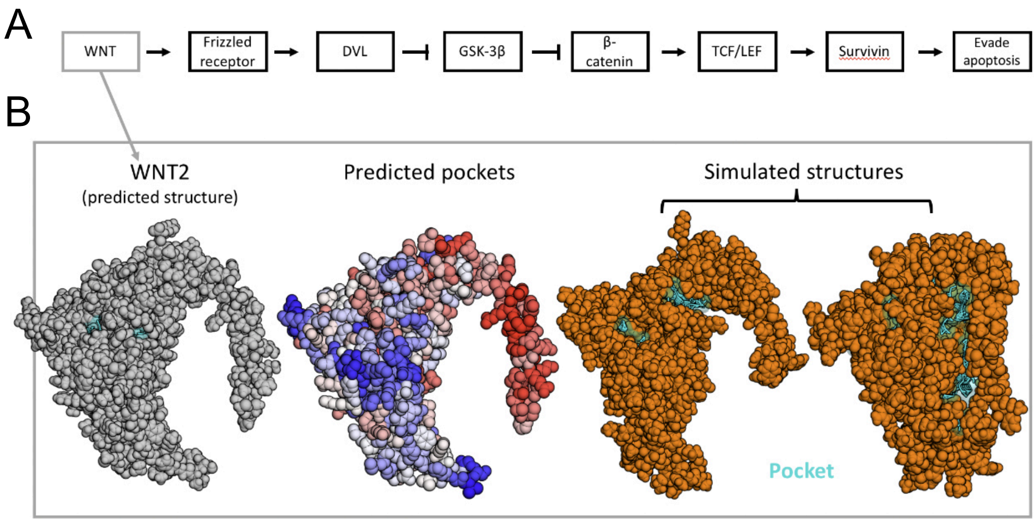


**Figure S8:** PocketMiner predicts a cryptic pocket in Wnt2 that opens in simulation. A) Wnt2 is part of the Wnt2 signal transduction pathway that regulates apoptosis and has been identified as a cancer target. B) PocketMiner predicts a cryptic pocket will form based on the AlphaFold-predicted structure of Wnt2. In simulation, a cryptic pocket forms as a result of an interdomain closure.

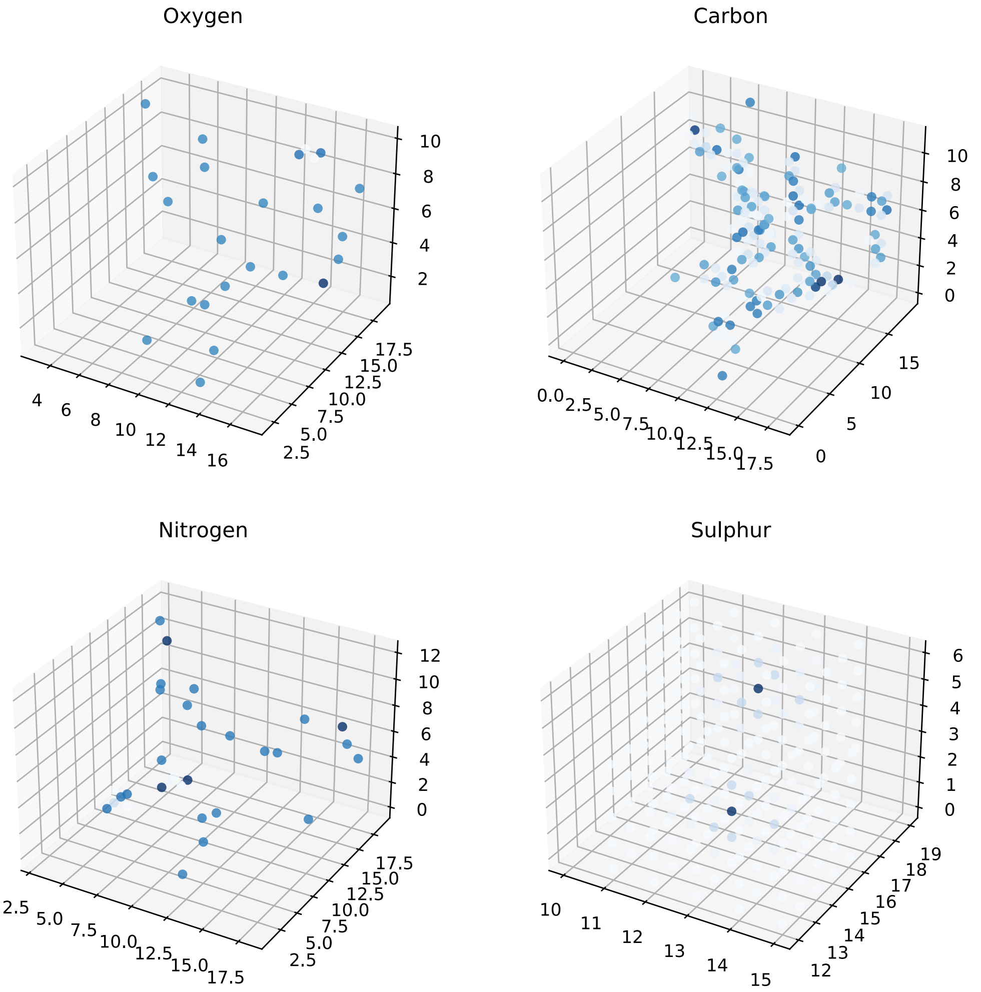


**Figure S9:** Schematic depicting 3D grid featurization for 3D-convolutional neural networks. There are four separate channels, one for each of the elements shown above.

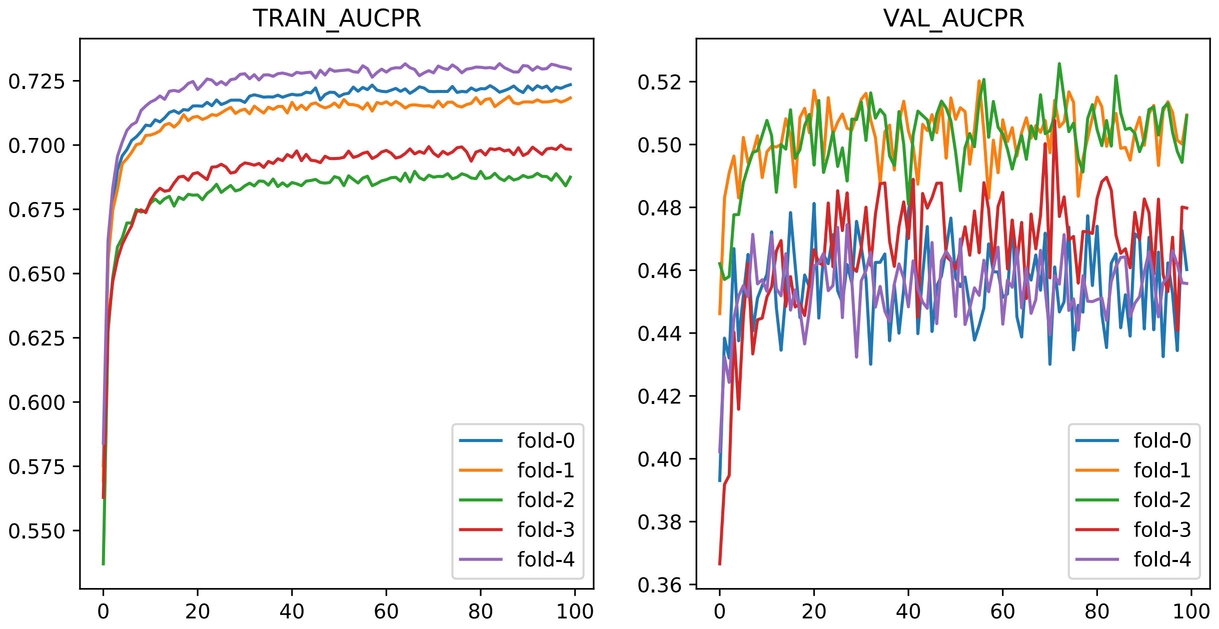


**Figure S10:** 3D convolutional neural network training and validation PR-AUC as a function of epoch (x-axis) demonstrate convergence around 20 epochs.

**Supplementary tables**

Contents

1. 5-fold cross validation network performance on task 2 validation set
2. Highest sequence identities among proteins in this work
3. Featurization performance
4. FAST simulations
5. PocketMiner and CryptoSite protein-level performance
6. PocketMiner sensitivity by protein topology and cryptic pocket type

**Table S1: 5-fold cross validation network performance on task 2 validation set.** We trained networks on sets of labels sampled from those generated in simulations as described under the parameters section. Intermediate labels were those with volume changes between 116 and 20 LIGSITE grid points.

The following weighting and sampling schemes were used:

- none:
- weighting:
- oversampling:
- undersampling:
- constant size balancing:

The positive label fraction across all labels is 0.1

| **Parameters** | | | | **Performance on task 2 validation set cryptic pocket examples** | | | |
|---|---|---|---|---|---|---|---|
| **intermediate labels included as negatives** | **training label weighting or sampling scheme** | **number of residues drawn** | **residue batch size** | **ROC AUC** | | **PR AUC** | |
| | | | | **mean** | **stdev** | **mean** | **stdev** |
| yes | none | n/a | 4 | 0.799 | 0.139 | 0.417 | 0.188 |
| no | constant size balancing | 160 | 4 | 0.783 | 0.151 | 0.398 | 0.18 |
| yes | undersampling | n/a | n/a | 0.786 | 0.144 | 0.392 | 0.174 |
| no | oversampling | n/a | n/a | 0.773 | 0.137 | 0.384 | 0.166 |
| no | weighting | n/a | 4 | 0.778 | 0.149 | 0.383 | 0.166 |

| | | | | | | | |
|---|---|---|---|---|---|---|---|
| no | none | n/a | 4 | 0.781 | 0.14 | 0.375 | 0.152 |
| yes | weighting | n/a | 32 | 0.772 | 0.13 | 0.374 | 0.163 |
| yes | constant size balancing | 160 | 4 | 0.76 | 0.158 | 0.365 | 0.175 |
| yes | weighting | n/a | 4 | 0.749 | 0.169 | 0.363 | 0.171 |
| no | constant size balancing | 160 | 32 | 0.751 | 0.149 | 0.361 | 0.172 |
| yes | constant size balancing | 160 | 32 | 0.752 | 0.154 | 0.355 | 0.162 |
| no | none | n/a | 32 | 0.764 | 0.132 | 0.354 | 0.152 |
| no | weighting | n/a | 32 | 0.744 | 0.145 | 0.34 | 0.158 |
| yes | none | n/a | n/a | 0.753 | 0.131 | 0.337 | 0.142 |
| yes | none | n/a | 32 | 0.744 | 0.141 | 0.329 | 0.143 |
| no | undersampling | n/a | n/a | 0.748 | 0.146 | 0.32 | 0.135 |
| no | none | n/a | n/a | 0.723 | 0.128 | 0.29 | 0.128 |
| yes | weighting | n/a | n/a | 0.733 | 0.129 | 0.29 | 0.122 |
| yes | oversampling | n/a | n/a | 0.729 | 0.136 | 0.288 | 0.127 |
| no | weighting | n/a | n/a | 0.723 | 0.138 | 0.282 | 0.136 |

**Table S2: Highest sequence identities among proteins in this work.** *Apo* PDB IDs are given where applicable, and the PBD IDs of SARS and MERS protein structures (or those used for homology modeling) can be found in the training_5_fold_cv tab of the attached SI spreadsheet. The complete list of sequence identities between proteins used in this study exceeding 40% can be found in the sequence_identity tab of the same spreadsheet.

| **Protein 1** | **Protein 2** | **Percent identity** | **Notes** |
|---|---|---|---|
| SARS-1-nsp16 | SARS-2-nsp16 | 93.493 | All in the same 5-fold cross validation fold |
| MERS-nsp16 | SARS-2-nsp16 | 65.870 | |
| MERS-nsp16 | SARS-1-nsp16 | 64.726 | |
| 1IGD | SARS-2-nsp12 | 59.016 | See fig S6A |
| 1IGD | SARS-2-nsp13 | 57.377 | See fig S6B |
| 2FD7 | 2OHG | 54.348 | See fig S6C |
| 1IGD | 1KMO | 52.459 | See fig S6D |

| | | | |
|---|---|---|---|
| 2FD7 | SARS-2-nsp12 | 52.174 | - |
| SARS-2-nsp12 | SARS-2-nsp7 | 50.633 | See fig S6E |
| 2FD7 | SARS-2-nsp13 | 50.000 | - |
| 1IGD | 2GG4 | 49.180 | - |
| 1JEJ | 2FD7 | 47.826 | - |
| 1V2N | 2FD7 | 47.826 | - |
| 5NZM | SARS-2-nsp7 | 46.835 | See fig S6F |

**Table S3: Featurization performance.** Performance of the featurization scheme used to generate training labels from the simulations above on the same 12 CryptoSite proteins on which it was optimized. Residues within 5Å of the cryptic site ligand in the *holo* crystal structure were used as positive true labels.

| | **PR AUC** | **ROC AUC** | **class split** |
|---|---|---|---|
| **4AKE** | 0.457 | 0.848 | 0.136 |
| **1BSQ** | 0.644 | 0.908 | 0.099 |
| **1EX6** | 0.409 | 0.838 | 0.091 |
| **1ALB** | 0.552 | 0.898 | 0.160 |
| **1NEP** | 0.771 | 0.903 | 0.185 |
| **1NI6** | 0.553 | 0.900 | 0.099 |
| **2BLS** | 0.384 | 0.908 | 0.025 |
| **2QFO** | 0.254 | 0.738 | 0.097 |
| **3F74** | 0.272 | 0.762 | 0.111 |
| **1EXM** | 0.117 | 0.708 | 0.074 |
| **1ADE** | 0.417 | 0.900 | 0.056 |
| **1MY0** | 0.504 | 0.923 | 0.070 |
| **mean ± 1 SD** | 0.445 ± 0.171 | 0.853 ± 0.072 | 0.100 ± 0.044 |

**Table S4: FAST simulations.** We ran the FAST adaptive sampling algorithm (see methods) on 15 proteins from CryptoSite, 9 highly rigid proteins, and 10 proteins with many *holo* crystal structures in order to generate training data and negative examples for our validation and test sets. The number of FAST rounds and RMSD cluster radius used for simulations of each protein are given in the table below. FAST was run with 10 40 ns long parallel simulations per round for all proteins.

| **Protein set** | **PDB ID** | **cluster radius** | **number of rounds of FAST** |
|---|---|---|---|
| Training | 1ADE | 0.14 | 5 |
| | 1ALB | 0.125 | 5 |
| | 1BSQ | 0.1 | 5 |
| | 1EX6 | 0.18 | 5 |
| | 1EXM | 0.14 | 7 |
| | 1MY0 | 0.14 | 5 |
| | 1NEP | 0.08 | 5 |
| | 1NI6 | 0.14 | 5 |
| | 1OFV | 0.11 | 5 |
| | 1QYS | 0.12 | 5 |
| | 1RHB | 0.12 | 5 |
| | 1RTC | 0.14 | 5 |
| | 2BLS | 0.12 | 5 |
| | 2CM2 | 0.12 | 5 |
| | 2QFO | 0.13 | 5 |
| | 3F74 | 0.12 | 5 |
| | 4AKE | 0.2 | 5 |
| | 5BVL | 0.11 | 5 |
| Highly rigid proteins providing negative examples for the test set | 1AMM | 0.1 | 3 |
| | 1IGD | 0.14 | 3 |
| | 2ALP | 0.1 | 3 |
| | 2FD7 | 0.1 | 3 |
| | 4HJK | 0.1 | 3 |
| | 4TQL | 0.14 | 3 |
| Proteins with many *holo* crystal structures providing negative examples for the test set | 1BTP | 0.1 | 3 |
| | 1HCL | 0.14 | 3 |
| | 2OSS | 0.15 | 3 |
| | 2ZHV | 0.15 | 3 |
| | 3GZ0 | 0.12 | 3 |

| | | | |
|---|---|---|---|
| | 4APE | 0.15 | 3 |
| | 5E4P | 0.11 | 3 |
| | 5NZ5 | 0.13 | 3 |
| | 5UOJ | 0.17 | 3 |
| | 5X8U | 0.12 | 3 |
| Structures from AlphaFold | Cyclin 1A | 0.1 | 3 |
| | PIM2 | 0.13 | 3 |
| | WNT2 | 0.26 | 3 |

**Table S5: PocketMiner and CryptoSite performance on individual proteins.** PocketMiner and CryptoSite were both run on the test set assembled in this work, and the accuracy of the predictions on each protein was calculated. The first set of proteins were known to form cryptic pockets and hence served as source of positive residues. The second set of proteins contained residues which were very unlikely to form cryptic pockets based on experimental and simulation data. Hence, this set of proteins, comprised of hyper-rigid proteins and proteins with extensive ligand co-crystal structures, was used a source of negative residues.

| type | *apo* PDB ID | CryptoSite accuracy | PocketMiner accuracy |
|---|---|---|---|
| Cryptic pocket | 6hb0 | 0.667 | 1.000 |
| | 2lao | 0.667 | 0.933 |
| | 1urp | 0.769 | 1.000 |
| | 6ypk | 0.875 | 0.417 |
| | 3ugk | 0.762 | 0.976 |
| | 5g1m | 0.684 | 0.789 |
| | 5nzm | 0.889 | 0.963 |
| | 4v38 | 0.913 | 1.000 |
| | 2w9t | 0.850 | 0.550 |
| | 2hq8 | 0.760 | 0.600 |
| | 4r72 | 0.625 | 0.750 |
| | 5za4 | 0.875 | 0.875 |
| | 2cey | 0.611 | 1.000 |
| | 1kmo | 0.773 | 0.909 |
| | 5nia | 0.963 | 0.926 |
| | 2fjy | 0.789 | 0.421 |
| | 3p53 | 0.688 | 1.000 |

| | | | |
|---|---|---|---|
| | 1kx9 | 0.781 | 0.000 |
| | 1tvq | 0.857 | 1.000 |
| | 2zku | 0.774 | 0.903 |
| | 3nx1 | 0.636 | 0.636 |
| | 4i92 | 0.842 | 0.947 |
| | 3qxw | 0.650 | 0.150 |
| | 5h9a | 0.955 | 1.000 |
| Highly rigid protein | 2fd7 | 1.000 | 0.978 |
| | 4tql | 0.572 | 1.000 |
| | 2alp | 0.697 | 0.787 |
| | 1amm | 0.770 | 0.865 |
| Proteins with many holo crystal structures | 4ape | 0.814 | 0.873 |
| | 5uoj | 0.755 | 0.717 |
| | 3gz0 | 0.686 | 0.863 |
| | 1btp | 0.802 | 0.676 |
| | 2oss | 0.563 | 0.850 |
| | 5zn5 | 0.754 | 0.836 |
| | 5x8u | 0.732 | 0.813 |
| | **mean** | **0.766** | **0.800** |

**Table S6:** PocketMiner sensitivity by protein topology and cryptic pocket type for test set proteins. Proteins used as sources of negative examples were not included. One protein with no assigned CATH code (PDB ID 6YPK) was excluded from the **CATH class** category and one protein with a large rearrangement (PDB ID 2FJY) was excluded from both the **Direction** and **Motion type** categories.

| | **Class** | **Mean** | **Standard deviation** |
|---|---|---|---|
| CATH class | 1 | 0.492 | 0.394 |
| | 2 | 0.803 | 0.315 |
| | 3 | 0.895 | 0.138 |
| Direction | forward | 0.677 | 0.347 |
| | reverse | 0.927 | 0.089 |
| Motion type | Secondary structure change | 0.653 | 0.358 |
| | Interdomain motion | 0.944 | 0.079 |

| | | | |
|---|---|---|---|
| | Secondary structure element motion | 0.632 | 0.394 |
| | Loop motion | 0.818 | 0.237 |
